## Supplemental figure and table for "Neural Geometry from Mixed Sensorimotor Selectivity for Predictive Sensorimotor Control"

**Supplement list**

Table S1. Neural datasets
Table S2. RNN neural dynamics similarity
Table S3. RNN perturbation results
Table S4. Alternative models

Figure S1. Characterization of hand movement
Figure S2. Three example neurons of monkey C
Figure S3. Three example neurons of monkey G
Figure S4. Three example neurons of monkey D
Figure S5. Single-neuron fitting results
Figure S6. Decoding results of extended datasets
Figure S7. Decoding results and neural states across epochs
Figure S8. dPCA and subspace projection
Figure S9. Neural trajectory during preparatory and peri-movement periods
Figure S10. Neural state of target-motion modulated M1 neurons from three monkeys
Figure S11. Target-motion modulation on hand-speed-filtered trials
Figure S12. Decoding results and weight distributions of RNNs

| No. | Monkey | Date | Target-motion conditions | n (Trials) | N (Neurons) | Hand trajectory |
|---|---|---|---|---|---|---|
| 1 | C | 20221022 | 0 °/s, ±120 °/s, ±240 °/s | 772 | 95 | Yes |
| 2 | C | 20221023 | 0 °/s, ±120 °/s, ±180 °/s, ±240 °/s, ±360 °/s | 1257 | 97 | Yes |
| 3 | C | 20221024 | 0 °/s, ±120 °/s, ±180 °/s, ±240 °/s, ±360 °/s | 1301 | 90 | Yes |
| 4 | C | 20221025 | 0 °/s, ±120 °/s, ±180 °/s, ±240 °/s, ±360 °/s | 1378 | 100 | No |
| 5 | C | 20221026 | 0 °/s, ±180 °/s, ±360 °/s | 801 | 86 | No |
| 6 | C | 20221105 | 0 °/s, ±120 °/s, ±240 °/s | 856 | 68 | No |
| 7 | C | 20221117 | 0 °/s, ±120 °/s, ±240 °/s | 802 | 58 | Yes |
| 8 | G | 20190913 | 0 °/s, ±120 °/s, ±240 °/s | 903 | 85 | Yes |
| 9 | G | 20190914 | 0 °/s, ±120 °/s, ±240 °/s | 855 | 96 | Yes |
| 10 | G | 20190915 | 0 °/s, ±120 °/s, ±240 °/s | 801 | 86 | Yes |
| 11 | G | 20190916 | 0 °/s, ±120 °/s, ±240 °/s | 782 | 123 | No |
| 12 | D | 20180606 | 0 °/s, ±120 °/s, ±240 °/s | 752 | 39 | No |
| 13 | D | 20180701 | 0 °/s, ±120 °/s, ±240 °/s | 458 | 39 | No |
| 14 | D | 20180804 | 0 °/s, ±120 °/s, ±240 °/s | 772 | 55 | No |
| 15 | D | 20180805 | 0 °/s, ±120 °/s, ±240 °/s | 587 | 45 | No |

**Table S1. Neural datasets**

This table lists the basic information of all involved datasets.

| Pairs | CC1 | CC2 | CC3 | Disparity |
|---|---|---|---|---|
| C-M1-MO vs. C-M1-TO | 0.84 | 0.80 | 0.64 | 0.88 |
| C-M1-MO-1 vs. C-M1-MO-2 | 0.95 | 0.90 | 0.85 | 0.63 |
| C-M1-MO vs. C-PMd-MO | 0.90 | 0.75 | 0.73 | 0.83 |
| C-M1-MO vs. G-M1-MO-1 | 0.93 | 0.87 | 0.79 | 0.71 |
| C-M1-MO vs. G-M1-MO-2 | 0.93 | 0.89 | 0.79 | 0.70 |
| G-M1-MO-1 vs. G-M1-MO-2 | 0.97 | 0.87 | 0.82 | 0.51 |
| C-M1-MO vs. RNN-MO | 0.95±0.00 | 0.93±0.00 | 0.79±0.02 | 0.88±0.01 |
| G-M1-MO-1 vs. RNN-MO | 0.97±0.00 | 0.89±0.00 | 0.86±0.01 | 0.82±0.01 |
| G-M1-MO-2 vs. RNN-MO | 0.98±0.00 | 0.91±0.00 | 0.87±0.01 | 0.82±0.01 |
| C-M1-MO vs. RNN-MO-shuffle | 0.15±0.00 | 0.10±0.00 | 0.09±0.00 | 1.00±0.00 |

**Table S2. RNN neural dynamics similarity**

In **Pairs** column, the datasets used for comparison are listed. C-M1-MO-1 represents the neural response in M1 of monkey C, at the movement onset (MO) (±100ms), the 1-300 trials, on 2022/10/22; C-M1-TO came from the same trials, but at the target on (TO) (0-200 ms); C-PMd-MO came from the same trials, but in PMd; C-M1-MO-2 came from the same session but the 301-600 trials. G-M1-MO-1 means the neural response in M1 of monkey G, at the MO, the 1-300 trials on 2022/9/14, while G-M2-MO-2 came from the 1-300 trials on 2022/9/15. RNN-MO means the node activity at the MO [-200 ms, 120 ms] and RNN-MO-shuffle came from the same trials but temporally shuffled. **CC1** refers to the Pearson correlation coefficient between the first canonical components of the paired data, while **CC2** and **CC3** are similar, representing the coefficient between the second and third components, respectively. **Disparity** is the disparity value obtained with Procrustes analysis. RNN-included results are shown as mean ± sd. across 100 network models.

| | Distance Error | $R^2$ of fitting ellipses |
|---|---|---|
| Intact models | 0.0046 ± 0.0027 | 0.9781 ± 0.0498 |
| **Ablated nodes** | | |
| S | 0.1358±0.0147 | 0.9351±0.0856 |
| G | 0.1438±0.0141 | 0.9487±0.0644 |
| A | 0.1533±0.0184 | 0.9510±0.0642 |
| S only | 0.0969±0.0278 | 0.9019±0.0904 |
| G only | 0.0752±0.0346 | 0.9448±0.0707 |
| A only | 0.1133±0.0335 | 0.9337±0.0797 |
| **Changed connections** | | |
| S → S | 0.0697 ± 0.0279 | 0.8361 ± 0.0975 |
| S → G | 0.0687 ± 0.0300 | 0.8891 ± 0.0956 |
| S → A | 0.0815 ± 0.0311 | 0.8473 ± 0.1140 |
| G → S | 0.0670 ± 0.0294 | 0.8499 ± 0.1051 |
| G → G | 0.0598 ± 0.0263 | 0.8646 ± 0.1161 |
| G → A | 0.0773 ± 0.0308 | 0.8321 ± 0.1139 |
| A → S | 0.0763 ± 0.0287 | 0.8415 ± 0.1063 |
| A → G | 0.0777 ± 0.0320 | 0.8440 ± 0.1198 |
| A → A | 0.0803 ± 0.0296 | 0.7818 ± 0.1265 |

**Table S3. RNN perturbation results**

S for PD shift, G for Gain modulation, and A for Addition. The **Ablated nodes** column shows the modulation type of which the nodes were ablated (all connection weights set to zero). The difference between S and S only is that the former includes nodes with mixed selectivity, while the latter includes nodes merely with PD-shift modulation. It's similar to G/A and G only/A only. In the **Changed connections** case, the connection structure of each network was perturbed by making the connection weight between certain modulation groups 1.5 times the original. S → S indicates the connection from S modulation nodes to S modulation nodes, with others like S → G being similar. This perturbation would worsen the network performance (top) and decrease the fitting goodness of conditional ellipses (bottom). Both of the performance and fitting influences show significant between-group differences (K-W test, $p<0.001$). All results are shown as mean ± sd. across 100 network models.

| Models | GM | GT | sINIT |
|---|---|---|---|
| Performance | 0.0052±0.0032 | 0.0130±0.0075 | 0.0131±0.0071 |
| Correction rate | 1.00±0.01 | 0.84±0.13 | 0.82±0.15 |
| Gain (G) nodes % | 51.9±5.7% | 68.0±6.1% | 50.8±12.9% |
| PD shift (S) nodes % | 18.7±6.2% | 56.1±3.3% | 48.7±15.0% |
| Addition (A) nodes % | 68.6±5.8% | 61.9±5.4% | 61.0±8.8% |
| No modulation nodes % | 12.3±5.4% | 7.1±3.5% | 10.7±6.9% |
| No activation nodes % | 40.6±3.9% | 62.1±31.0% | 70.7±24.0% |
| PC1 explained variance % | 54.2±3.0% | 38.1±2.3% | 52.6±6.9% |
| PC2 explained variance % | 42.4±2.3% | 28.1±2.8% | 32.1±5.9% |
| PC3 explained variance % | 2.4±1.0% | 14.3±2.6% | 7.1±1.5% |
| $R^2$ of fitting ellipses | 0.9664±0.0281 | 0.7616±0.1655 | 0.8507±0.0687 |
| $R^2$ of fitting ellipses (Dim 1) | 0.9909±0.0173 | 0.8365±0.1365 | 0.8905±0.0512 |
| $R^2$ of fitting ellipses (Dim 2) | 0.9880±0.0214 | 0.7857±0.2105 | 0.8781±0.0683 |
| $R^2$ of fitting ellipses (Dim 3) | 0.0170±0.0151 | 0.4314±0.3281 | 0.3722±0.3400 |
| $R^2$ of fitting tilting angle | 0.0062 | 0.2528 | 0.4614 |

**Table S4. Alternative models**

Here GM represents the network models only receiving a GO-signal and motor intention; GT represents the network models only receiving a GO-signal and target location; sINIT represents the network models whose hidden connection weights were initialized in sparse distribution. All results, except for the R-squares of fitting tilting angle, are presented as mean ± sd. across 10 network models for each setting, while the R-squares of fitting tilting angle were calculated from pooled neural state points.

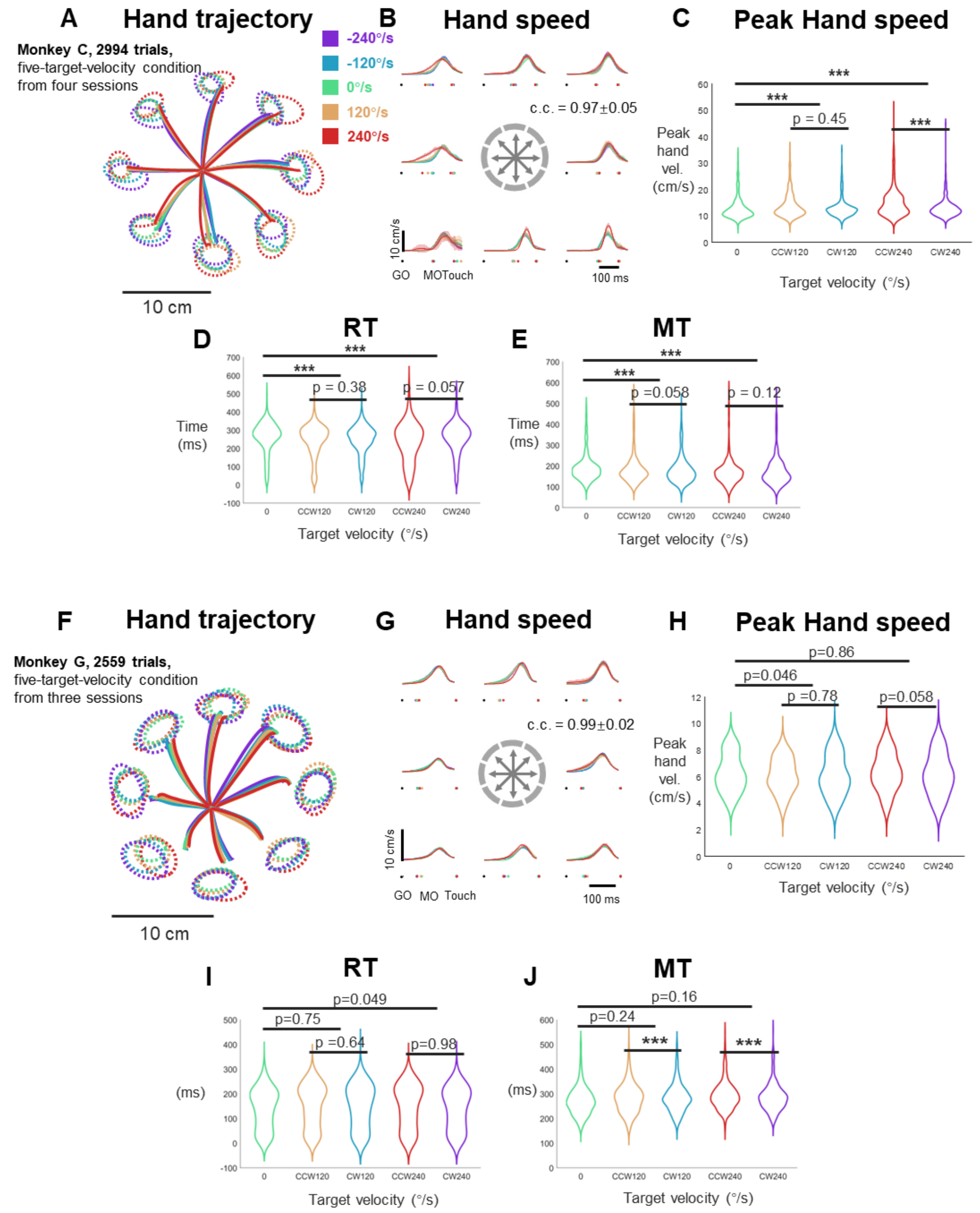


**Figure S1. Characterization of hand movement**

**A** The trial-averaged hand trajectory and touch endpoint distribution were averaged in five target-velocity and eight reach-direction conditions. All trajectories were colored according to the target-motion conditions, including monkey C's 2994 successful trials from four sessions.

**B** The trial-averaged hand speed was calculated from hand trajectory data (same in **A**), aligned to the GO (black dots), the averaged MO and the Touch, in each target-motion condition (colors with same meaning to **A**). Single-trial MO is defined as the moment when hand velocity first rose to 5% of the maximum. The correlation coefficient of hand speed profiles is 0.97 ± 0.05, mean ± std. of 40 conditions.

**C** The distribution of peak hand speed in five target-motion conditions (same as in **B**). The peak hand speed in the static-target condition was smaller than that in the ±120°/s and ±240°/s conditions (ANOVA p-value of 0°/s vs. ±120°/s, ±120°/s vs. ±240°/s were $<10^{-6}$, $<10^{-20}$). The hand speed of CCW and CW conditions differed little (ANOVA p-value of CCW vs. CW was 0.45 within 120°/s and $10^{-4}$ within 240°/s).

**D**-**E** The distribution of reaction time (RT) and movement time (MT) in five target-motion conditions (same sessions as in **A**-**C**). The temporal durations of the static-target condition were larger than 120°/s and 240°/s conditions (ANOVA p-value of 0°/s vs. ± 120°/s, ±120°/s vs. ±240°/s were $<10^{-8}$ and $<10^{-9}$ for RT, $<10^{-4}$ and $<10^{-12}$ for MT). The difference between durations of CCW and CW conditions were insignificant (ANOVA test).

**F-G** The trial-averaged hand trajectory and speed of monkey G for 2559 successful trials from four sessions.

**H-J** The distribution of peak hand speed, RT, and MT in five target-motion conditions (same sessions from monkey G; three stars mean ANOVA test $p<0.01$).

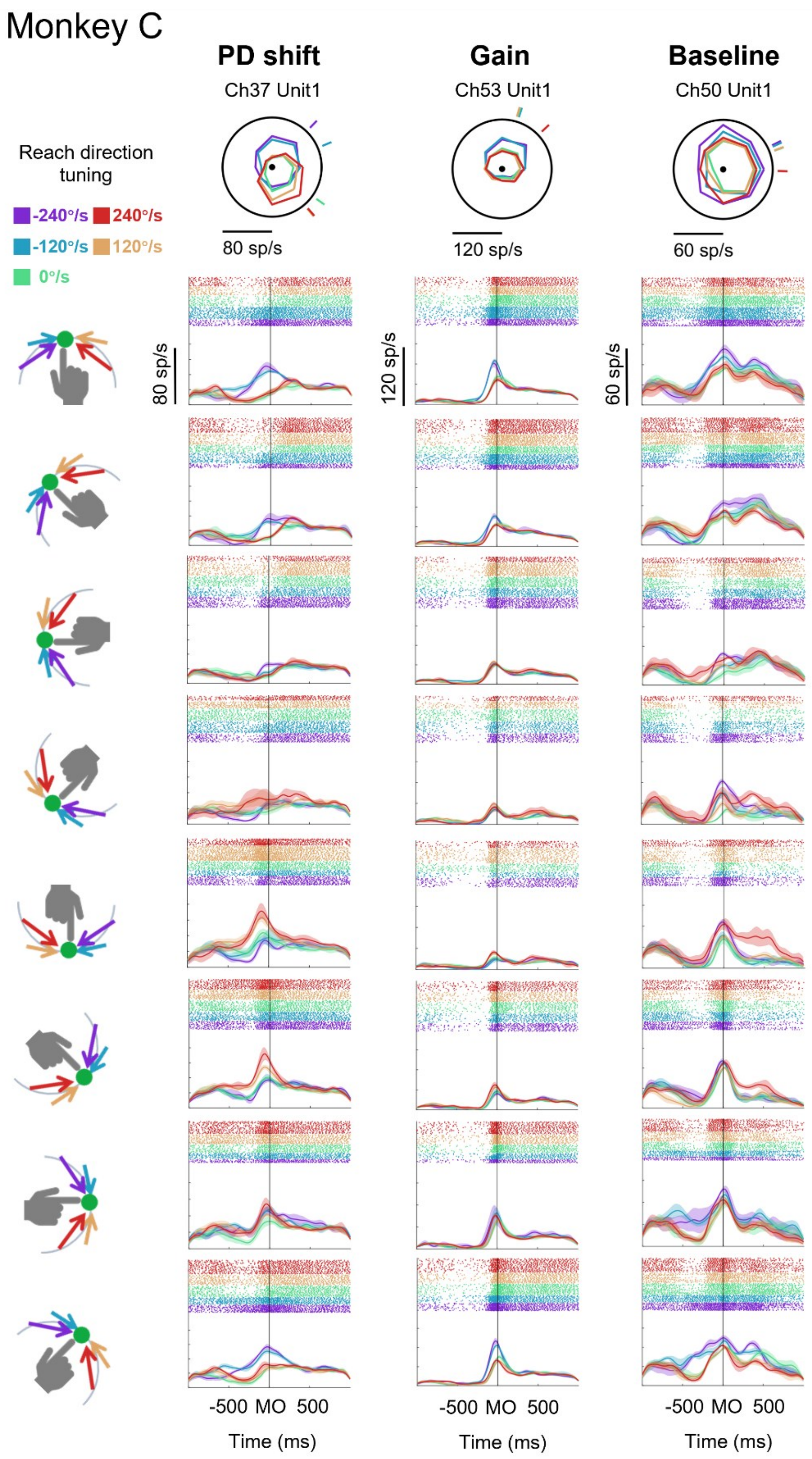


**Figure S2. Three example neurons of monkey C**

From left to right, each column shows one example neuron with labeled modulation. In the first row, the radar subplots show the neuronal reach-direction tuning curves at the movement onset (MO ± 0.1 s) in five target-motion conditions, with short lines outside the circle marking corresponding preferred directions. The next eight rows show rasters and peristimulus time histograms (PSTHs) of the example neuron in eight reaching areas (as indicated on the left, averaged trials were with touch endpoints within the 45° sector), aligned to the MO, in five target-motion conditions. In all subplots, the colors represent the corresponding target-motion conditions as shown by legends at the left side of the first row.

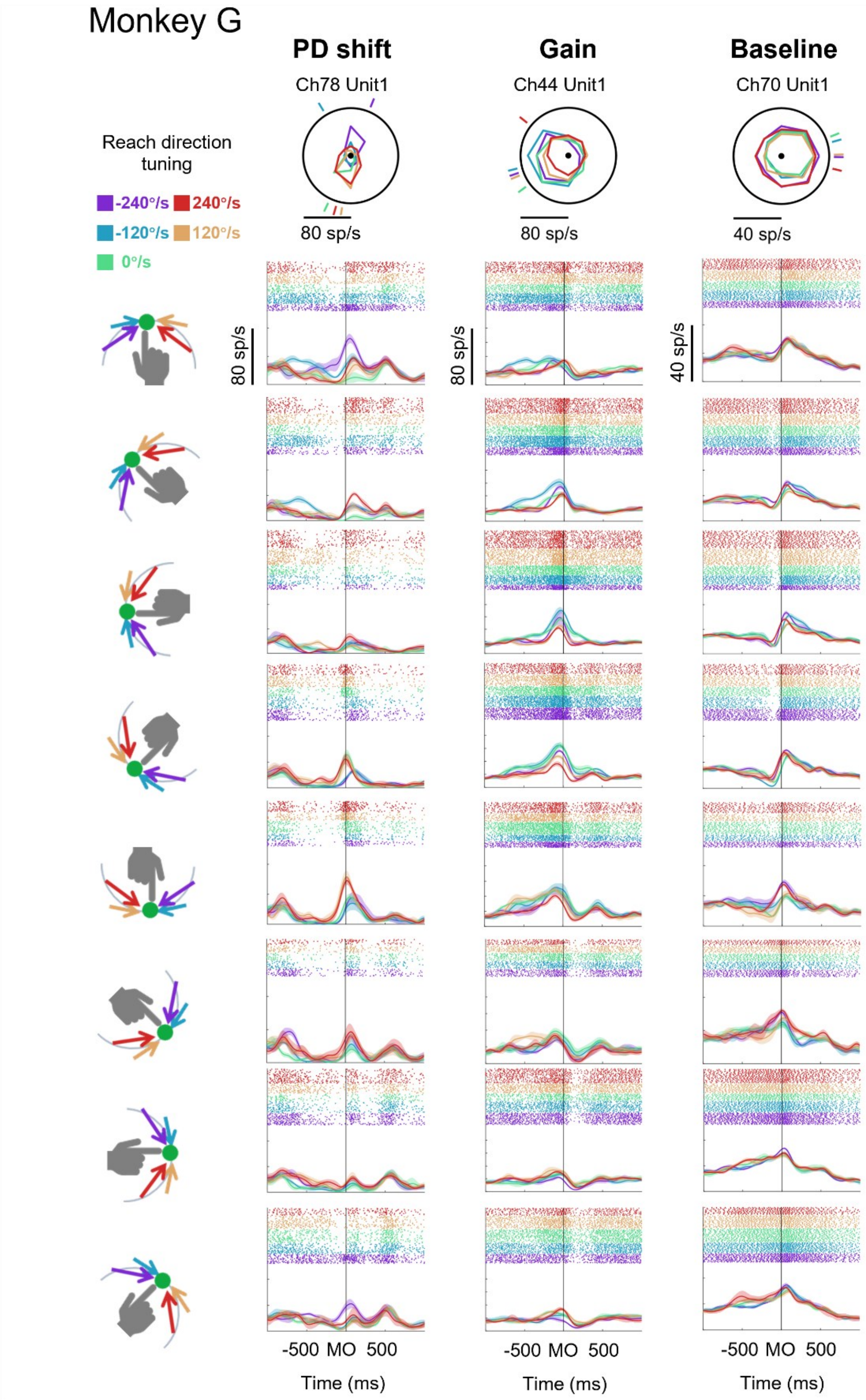


**Figure S3. Three example neurons of monkey G**

Legends are similar with Figure S2.

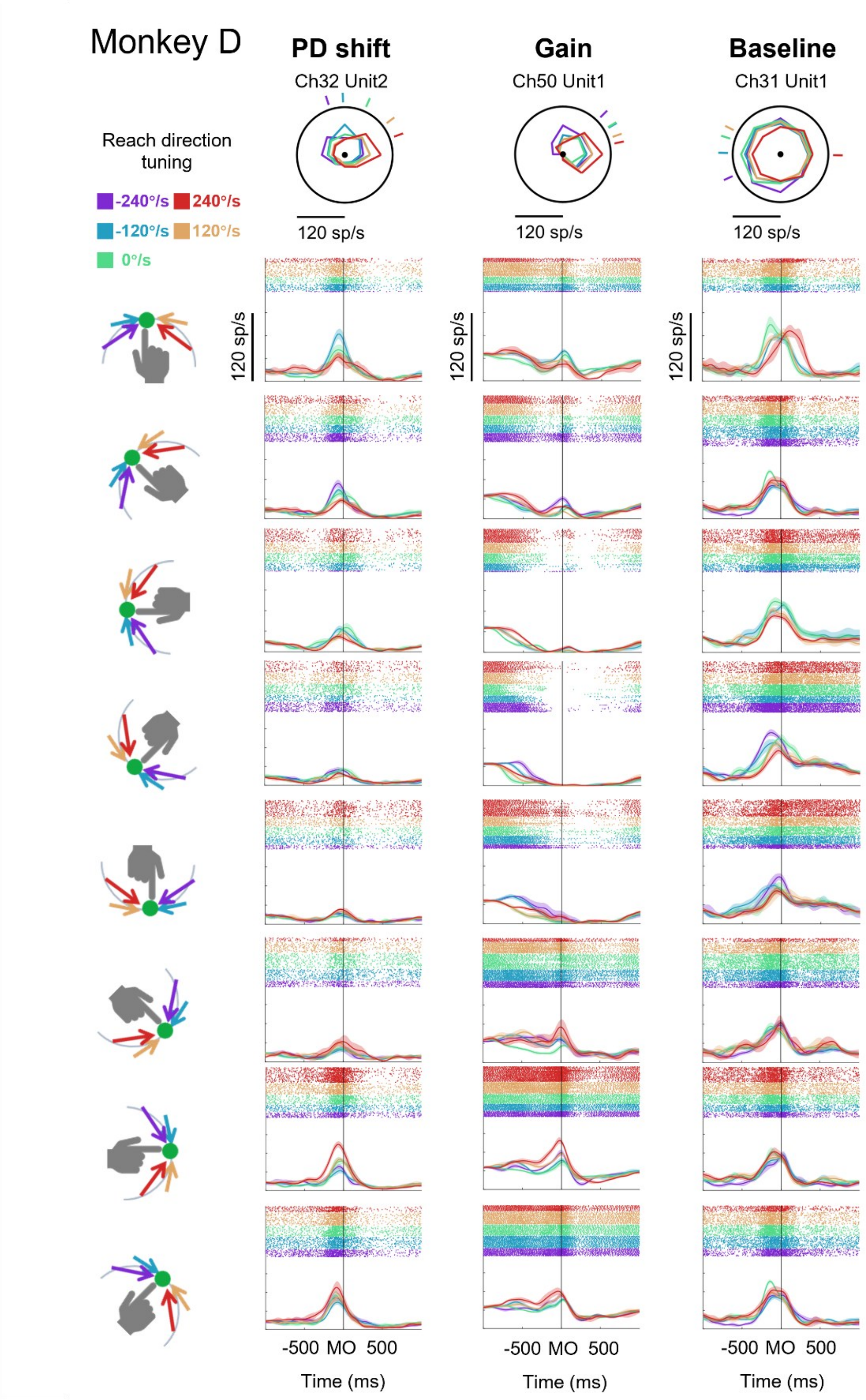


**Figure S4. Three example neurons of monkey D**

Legends are similar with Figure S2.

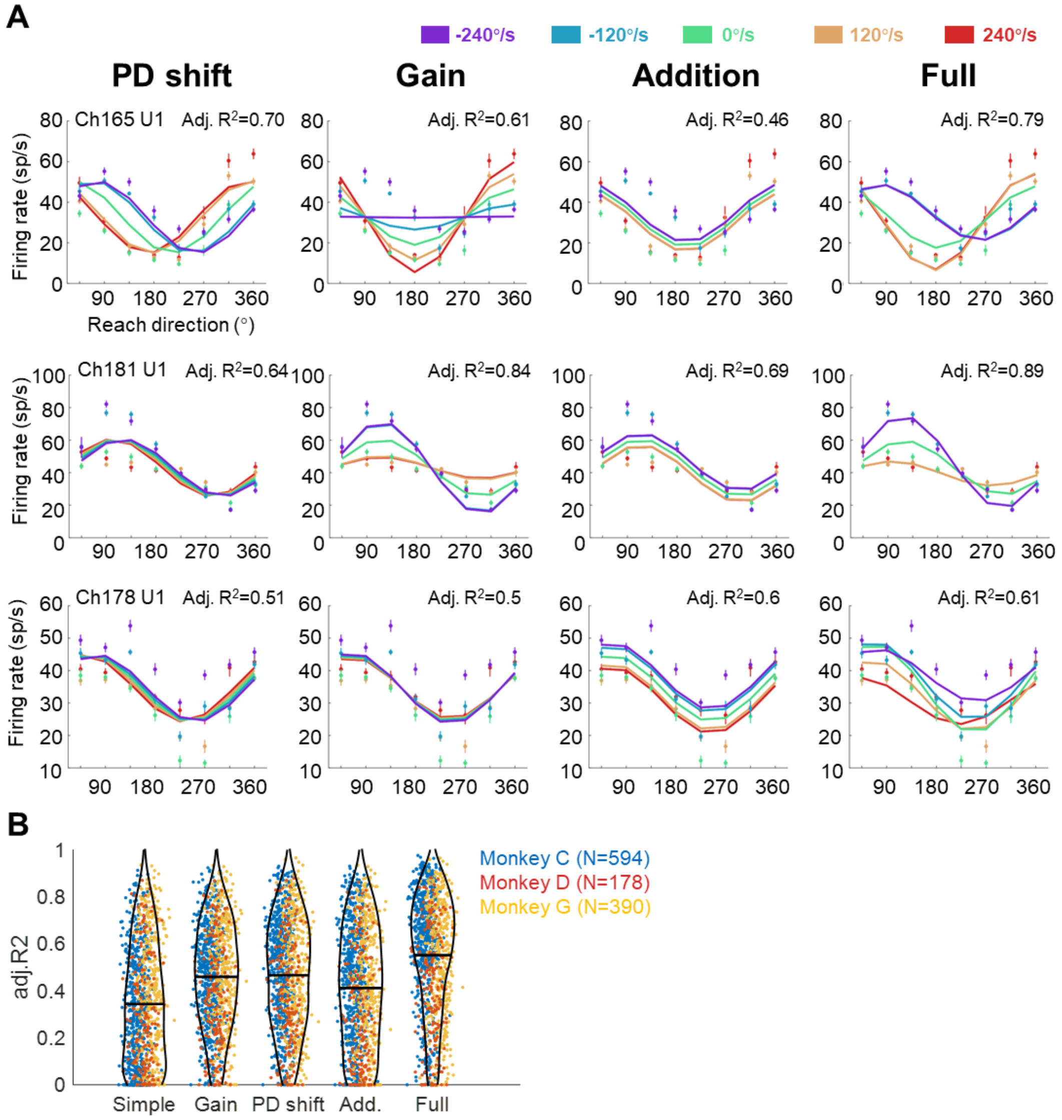


**Figure S5. Single-neuron fitting results**

**A** Three example neuron was fitted with the PD shift, gain, baseline, and full models. Each row, the subplots show the fitting results of one example neuron. Dots and bars indicate the mean and standard deviation of condition-averaged neuronal activity. Solid lines represent fitted tuning curves. The adjusted R-square is labelled at the top right corner of each subplot.

**B** The comparison of fitting goodness among five models (above four and a simple model only fitting reach direction with cosine function). Each dot shows the adjusted R-square of a single neuron (monkey C, N=594; monkey D, N=178; monkey G, N=390; in blue, red and yellow, respectively). The black bar is the mean of adjusted R-square (N=1162, simple model: 0.34±0.25, gain model: 0.46±0.22, PD shift model: 0.47±0.24, Addition model: 0.41 ±0.26, full model: 0.55±0.24, mean±sd.). The adjusted R-squares of the full model are obviously larger than the other models (Wilcoxon signed rank test, one-tailed, p<0.01).

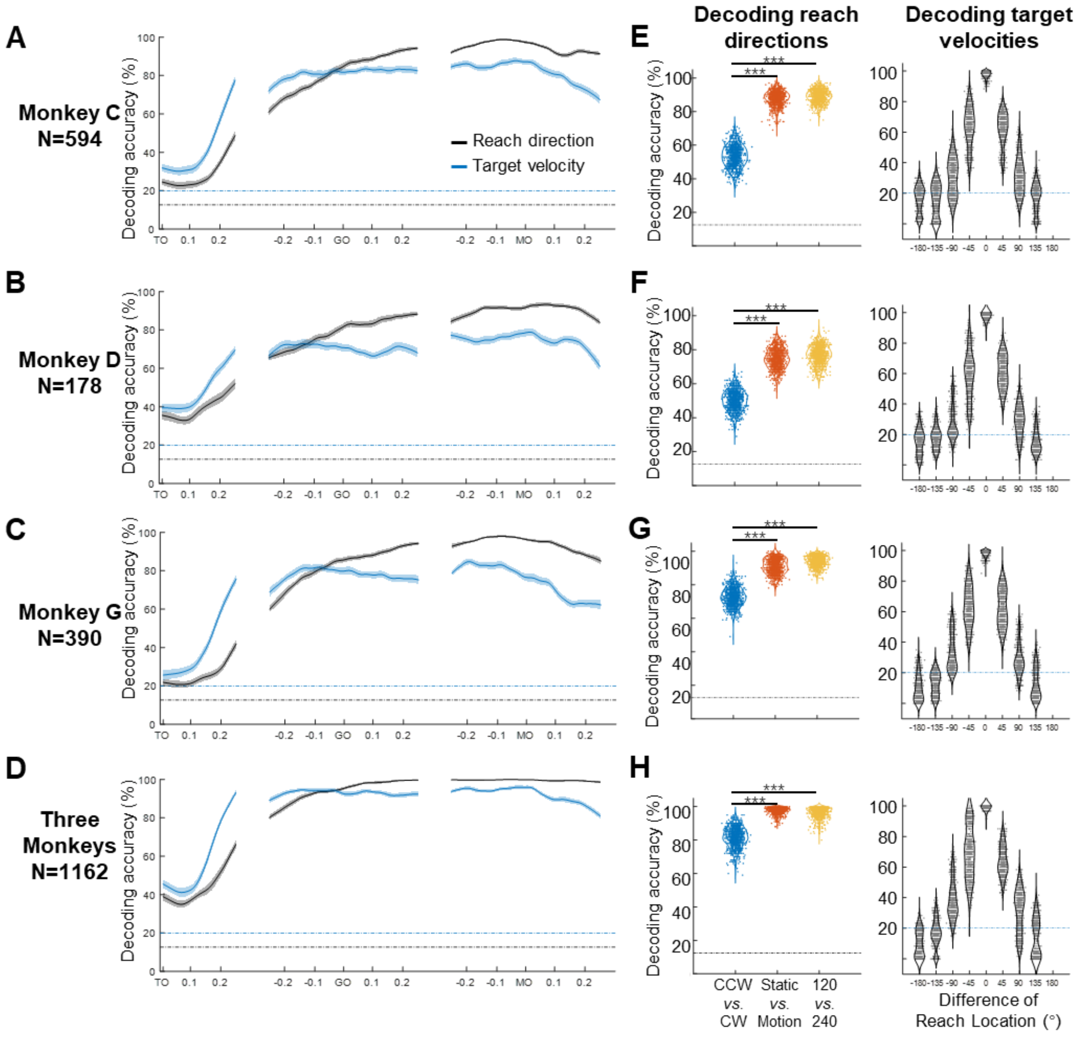


**Figure S6. Decoding results of extended datasets**

**A-D** The temporal decoding accuracy (SVM with 10-fold cross-validation) of reach direction (black line) and target velocity (blue line) on pseudo-population activity (merged Monkey C, D, G, and all monkeys), aligned to the target on (TO), GO, and movement onset (MO). We randomly selected the same number of trials to merge dataset, 40 condition by 15 repetitions without replacement for 600 trials, then trained and tested on the merged datasets. The dash and dotted lines are the chance levels of decoding reach direction (black, one in eight) and target velocity (blue, one in five). The shaded area indicates the standard deviation of the decoding accuracy for 10 repetitions.

**E-H** The left panel shows the performance of reach-direction decoder (chance level: one in eight) transferred between different target-motion conditions. The SVM decoder was built on 100 randomly selected trials from the training dataset, and tested in another 100 trials from a dataset of different conditions (CCW vs. CW, static vs. motion, 120 vs. 240). The distributions of decoding accuracy were collected from 1000 repetitions, and were compared with one tailed t-tests ($p<0.01$, with three stars). The right panel shows the performance of the target-velocity decoder (chance level: one in five) in different reach-direction conditions. In this case, reach direction is grouped into eight equal sectors (each 45°), and for each condition 60 trials were randomly selected for training and another randomly selected 60 trials for testing. The accuracy distribution was also obtained with 1000 repetitions.

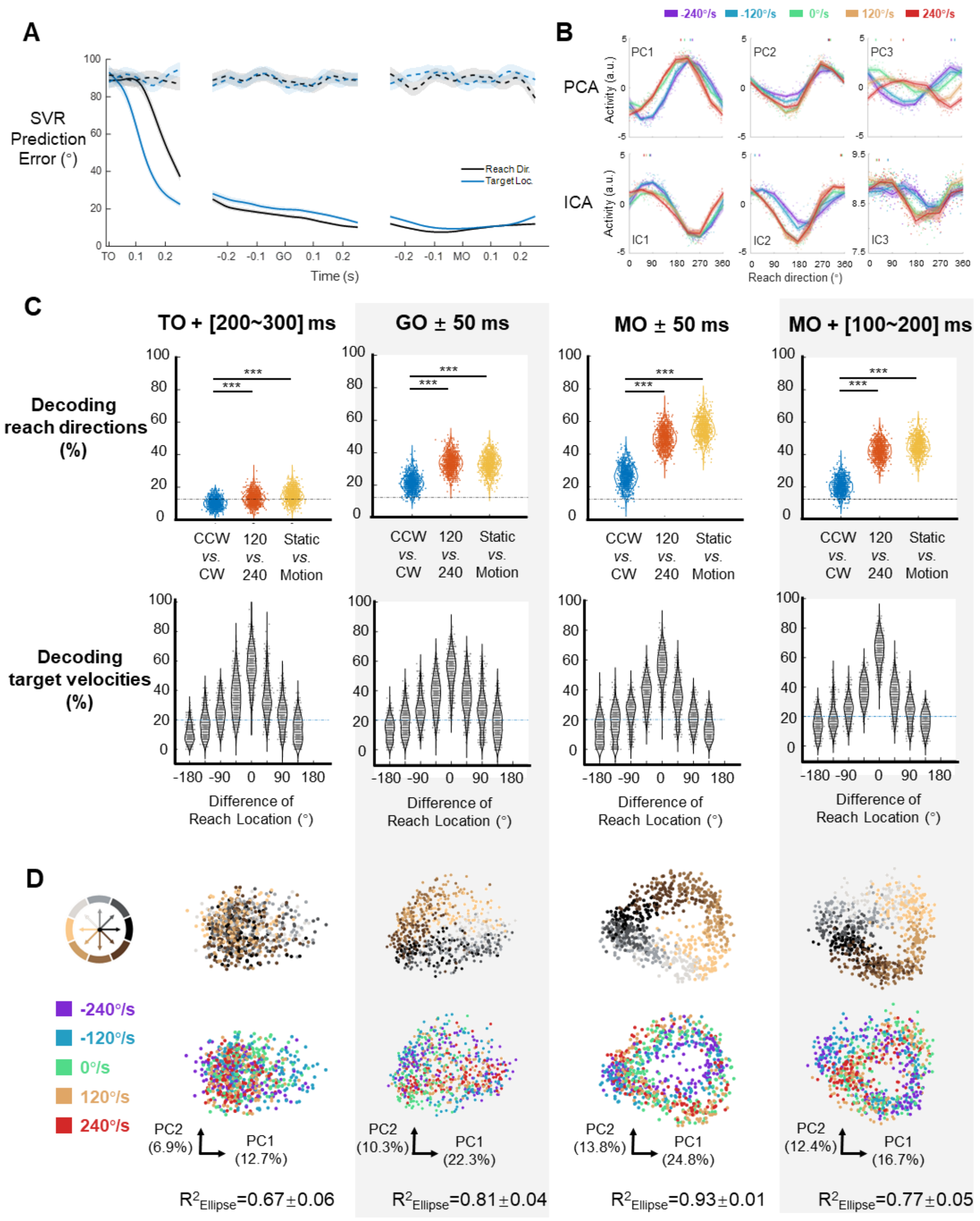


**Figure S7. Decoding results and neural states across epochs**

**A** The decoding accuracy (support vector regression, SVR with 10-fold cross-validation) of reach direction (black line) and real-time target direction (blue line) on population activity (monkey C, n=95, 772 trials), is aligned to the target on (TO), GO, and movement onset (MO). The dotted lines represent the chance level of decoding reach direction (black) and target direction (blue), using data with shuffled trial labels. The line and shaded area indicate the mean and standard error of the decoding accuracy for trials.

**B** Results of unsupervised dimensionality reduction on the example dataset. Each subplot shows the directional tuning of one component at the movement onset (PCAs are the same with **Figure.2D**). Each dot represents a trial, and tuning curves are averaged by eight reach directions and colored by target-motion conditions. The short lines at the top of the subplot show corresponding PDs by a weighted sum of response.

**C** Verifying the generalization of the decoder under different conditions with data from multiple time periods. The first row shows the performance of reach-direction decoder (chance level: one in eight) transferred between different target-motion conditions. The SVM decoder was built on randomly selected 100 trials in training dataset and tested in another 100 trials from a dataset of different conditions (CCW *vs.* CW, 120 *vs.* 240, static *vs.* motion). The distributions of decoding accuracy were from 1000 repetitions and compared with one tailed t-test ($p<0.01$, with three stars). The second row shows the performance of target-velocity decoder (chance level: one in five) transferred in different reach-direction conditions. In this case, reach directions are grouped by eight equal sectors (each 45°), and for each condition 60 trials were randomly selected for training and another 60 for testing. The accuracy distribution was obtained from 1000 repetitions.

**D** The neural states across epochs. In these planes spanned by the first two PCs, the dots represent the single-trial neural states in color corresponding to reach directions (first row) or target velocity (second row). The explained variance ratio is labelled on the corresponding axes. The goodness of fitting ellipses ($R^2_{Ellipse}$) for the state dots is shown as mean ± sd. across five target-motion conditions.

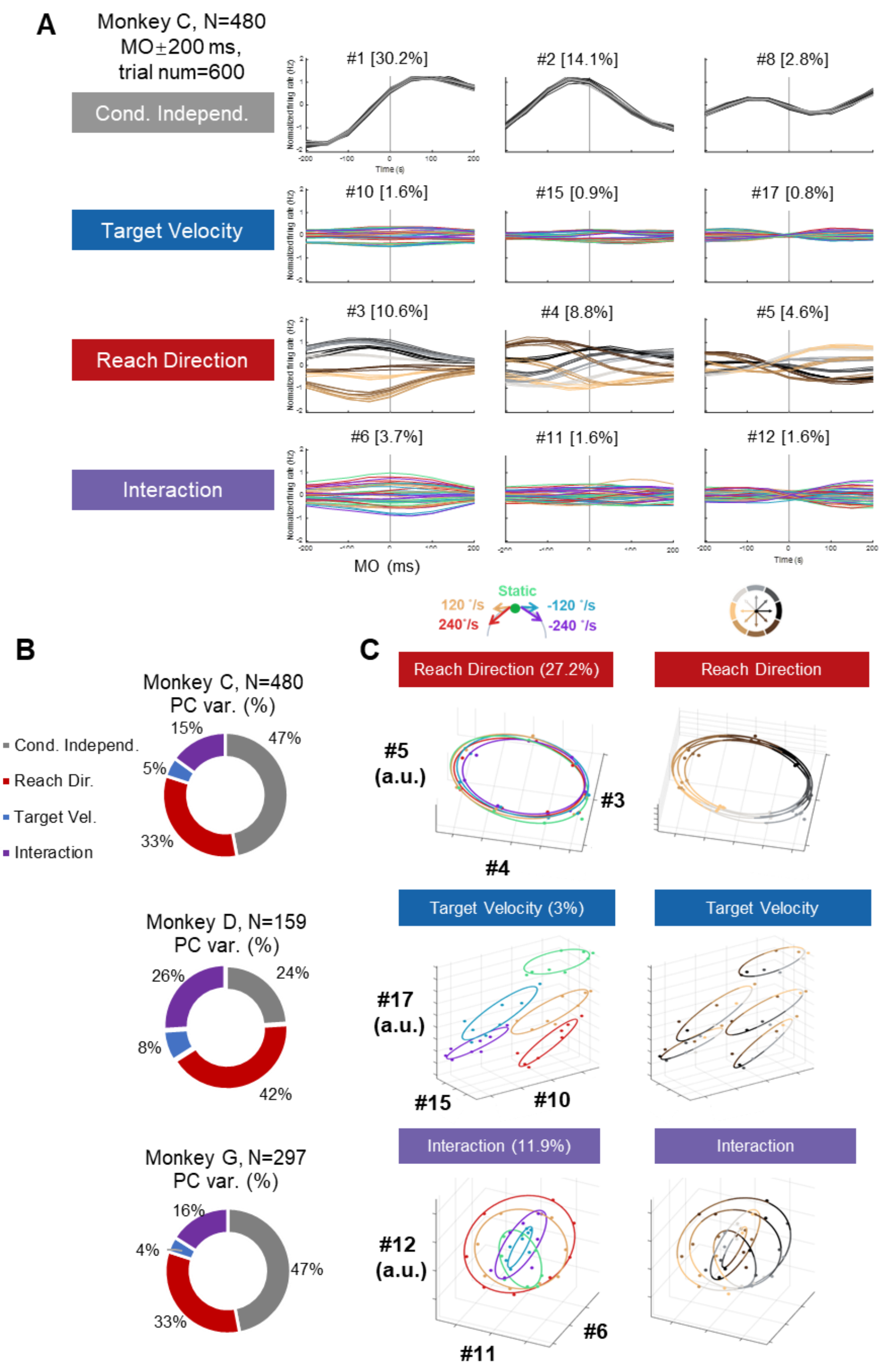


**Figure S8 dPCA and subspace projection**

**A** Neural data was merged to include target-velocity modulated units (480 selected neurons, 600 trials, eight time-bins with each 50 ms, from six sessions of monkey C). The firing rates were averaged by 40 conditions to perform dPCA. The condition-independent and reach direction components are colored according to reach directions, while the target velocity and interaction components are colored based on target-motion conditions.

**B** Summary of explained variance ratios of different dPCA components for the merged datasets of three monkeys.

**C** Neural states in “reach-direction subspace”, “target-velocity subspace”, and “interception subspace”. We used dPCA components with different features to construct three subspaces (same data in **A**, reach-direction space #3, #4, #5; target-velocity space #10, #15, #17; interaction space #6, #11, #12), and we projected trial-averaged data into these orthogonal subspaces using different colormaps. This approach allowed us to obtain a “potent subspace” coding reach direction and a “null space” for target velocity. The results showed that the reach-direction subspace effectively represented the reach direction. However, while the target-velocity subspace encoded the target velocity information, it still contained reach-direction clusters within each target-velocity condition, corroborating the results of the addition model in the main text (Figure 4). The interaction subspace revealed that multiple reach-direction rings were nested within each other, similar to the findings from the gain model (Figure 3 & 4). The interaction subspace also captured more variance than target-velocity subspace, consistent with our PCA results, suggesting the target-velocity modulation primarily coexists with reach-direction coding. Furthermore, we explored alternative methods to verify whether orthogonal subspaces could effectively separate the reach direction and target velocity. We could easily identify the reach-direction subspace, but its orthogonal subspace was relatively large, and the target-velocity information exhibited only small variance, making it difficult to isolate a subspace that purely encodes target velocity.

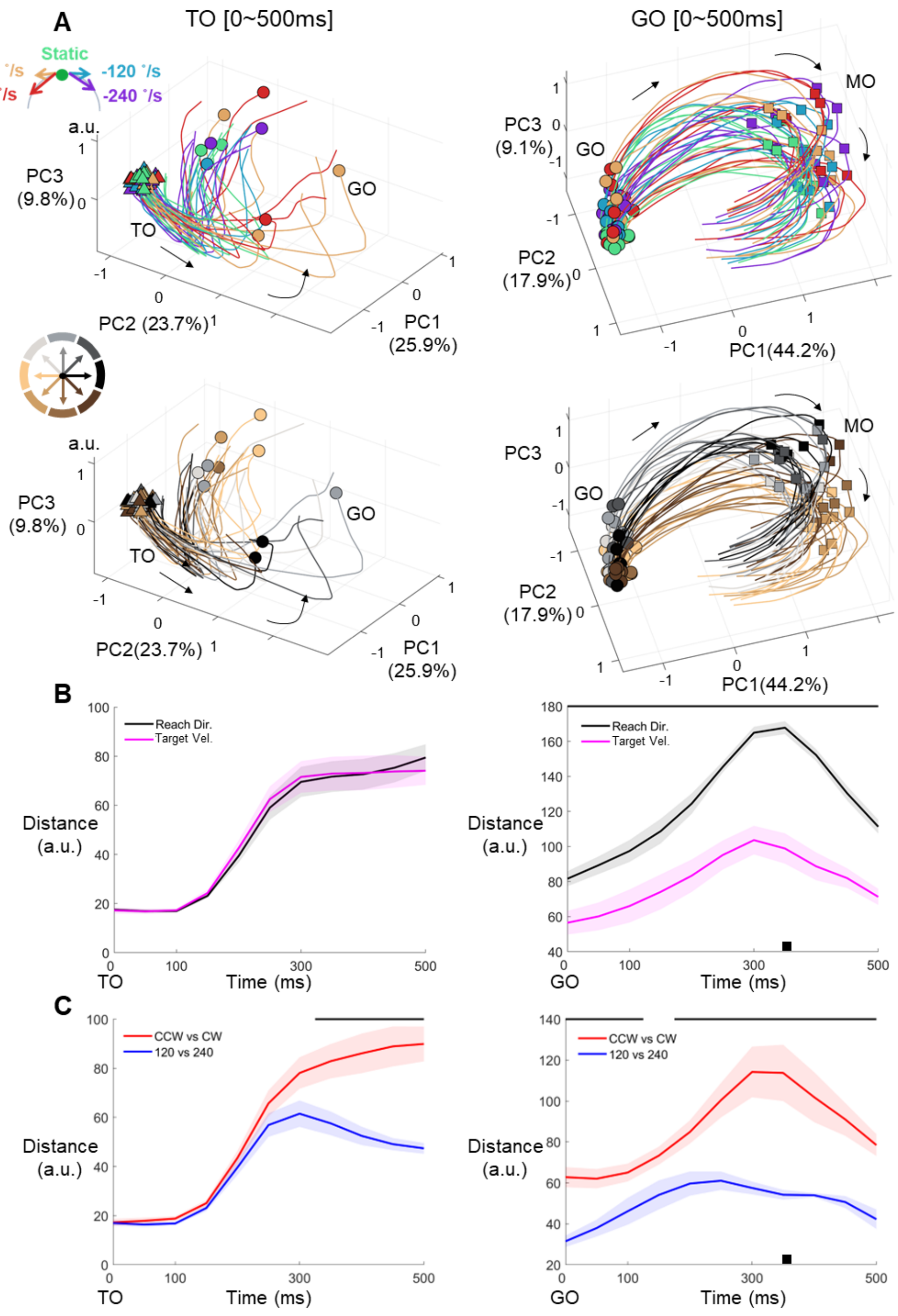


**Figure S9. Neural trajectory during preparatory and peri-movement periods**

**A** The neural trajectories (Monkey C, N=95, 40 conditions) in the subspace spanned by the first three PCs in two periods. Left column is delay period (from TO to TO+500 ms, TO means target onset), with triangles and circles marking condition-averaged TO and GO, respectively. Right column is the peri-movement period (from GO to GO+500 ms), with circles and squares marking GO and movement onset (MO), respectively. The same neural trajectories are plotted in two colormaps corresponding to target-motion conditions (first row) and reach directions (second row). Black arrows mark the direction of temporal evolution.

**B** The mean Euclidian distance of neural trajectories between each paired target-motion $c_5^2$ or reach-direction conditions $c_8^2$, aligned to TO (left) and GO (right). For target-motion conditions, the first eight PCs were included, while 14 PCs for reach-direction conditions. In both cases, the included PCs cover 90% explained variance. The black bar above the subplot marks those temporal bins in which there are significant differences between two distances (Kruskal-Wallis test, $p<0.05$). The shadow gives the standard error. The black square is the condition-averaged MO.

**C** Similar to **B**, but comparing CCW vs. CW and 120 vs. 240 conditions.

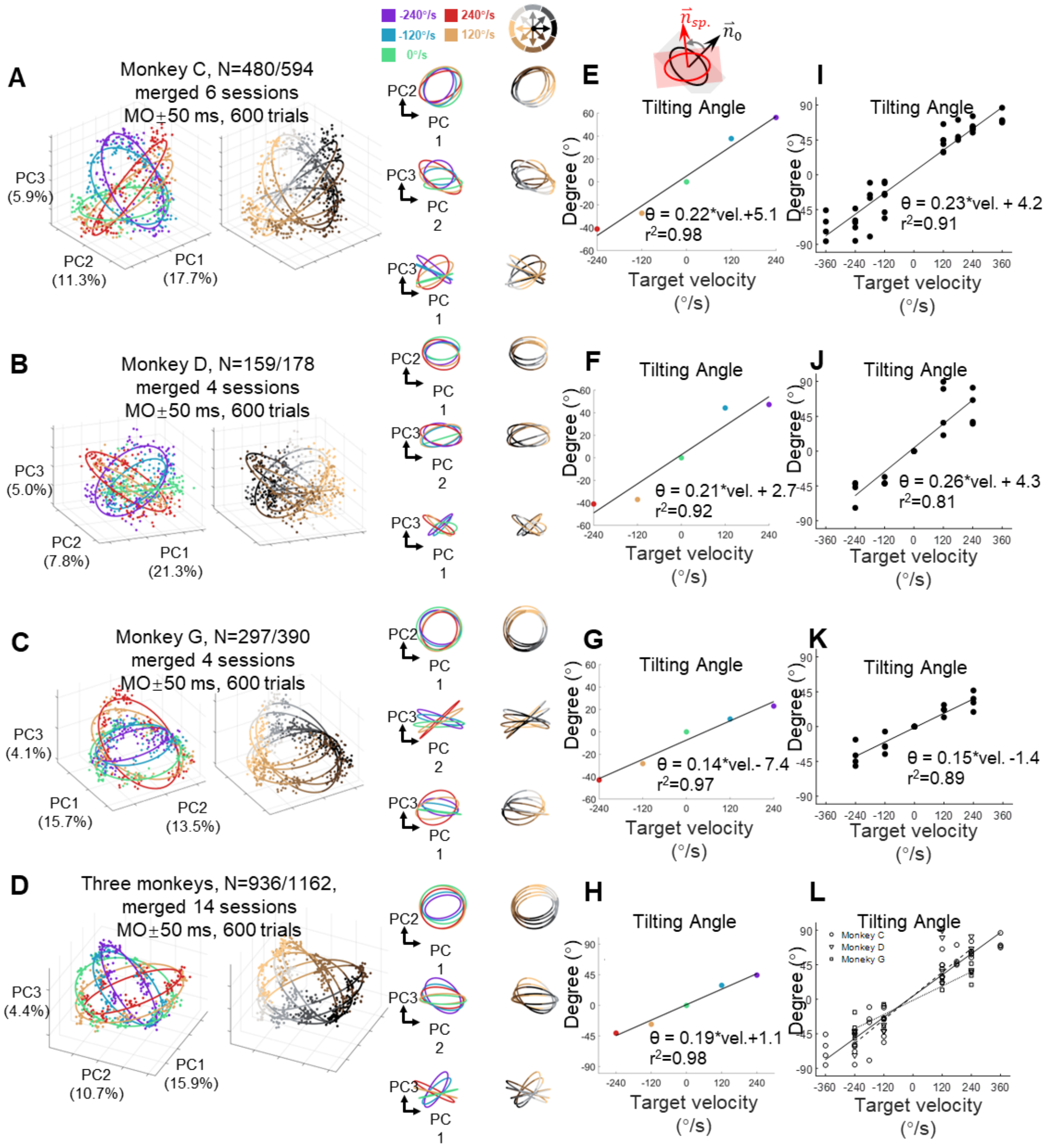


**Figure S10. Neural state of target-motion modulated M1 neurons from three monkeys**

**A-D** The neural state around the movement onset (MO ± 50 ms) was obtained for neurons selected from three monkeys. Neural data were collected from target-velocity modulated units (N) and randomly selected 15 trials in each of 40 conditions (K=600 trials). Each point represents the neural state of a single trial. Ellipses were fitted in each target-velocity condition. The legends can refer to Figure 3.

**E-H** The relative titling angle of ellipses from the static condition (merged datasets in **A-D**) has a linear relationship with target velocity. The five dots correspond to the tilting angles of five target-motion conditions.

**I-L** The titling angles for different datasets (as the same as in **A-D**).

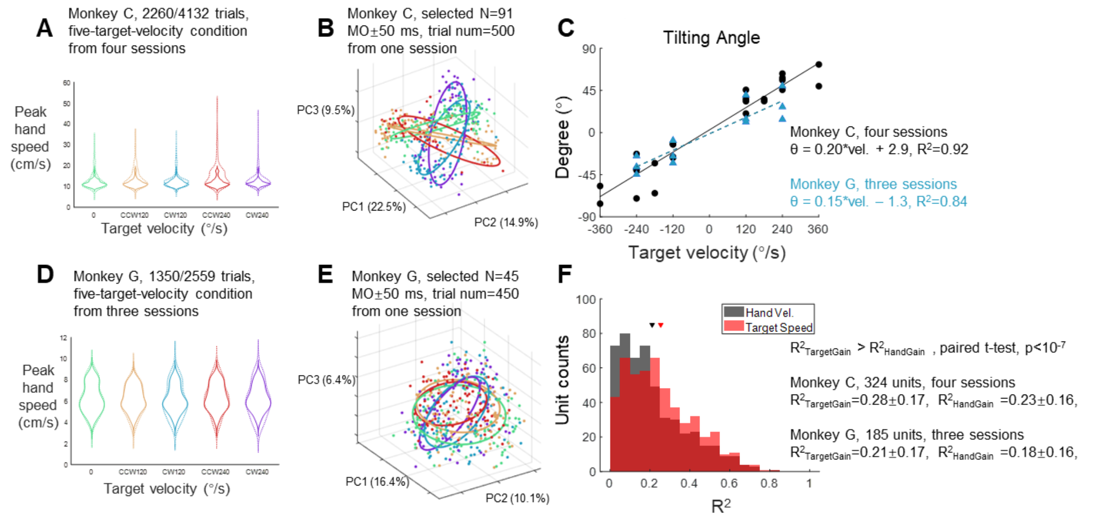


**Figure S11. Target-motion modulation on hand-speed-filtered trials**

**A** The distribution of peak hand speed of selected trials for monkey C. We used the 10th to 90th percentiles of peak hand speed in the static-target condition as the threshold range to select trials in the moving-target conditions. Solid lines indicate the new distribution (monkey C 2260/4132 trials), while dash lines show the original distribution from Figure S1C. After filtering, there is no significant difference between the distribution of peak hand speed in five target-motion conditions (ANOVA p-value of 0°/s vs. ±120°/s, 0°/s vs. ±240°/s, and ±120°/s vs. ±240°/s were 0.56, 0.14, and 0.26).

**B** Adjusted neural state of one example session from monkey C. In this session, 500 filtered trials (100 trials with peak hand speed within the threshold range were randomly selected for each target-motion condition), and 91 target-velocity modulated neurons are included.

**C** Tilting angle between ellipses of adjusted neural state. These angles were highly correlated with target velocity (black circles represent the neural state points from four sessions of monkey C, and blue triangles indicate those points from three sessions of monkey G).

**D** The distribution of peak hand speed of selected trials for monkey G. Similar to **A** (ANOVA p-value of 0°/s vs. ±120°/s, 0°/s vs. ±240°/s, and ±120°/s vs. ±240°/s were 0.60, 0.64, and 0.21).

**E** Adjusted neural state of one example session from monkey G, similar to **B**.

**F** The distribution of R-square for two single-neuron gain models. Both of the gain models can be expressed as $fr = c_0 + a*(y+c_1)*cos(x-b) + c_2*y$; where fr is the mean firing rate of MO±50 ms, x is reach direction. They only differ in the meaning of y, either target velocity or hand speed. The triangles indicate the mean R-square of the corresponding models, each marked with the same color as in the histogram. The gain model of target velocity performed better than that of hand speed.

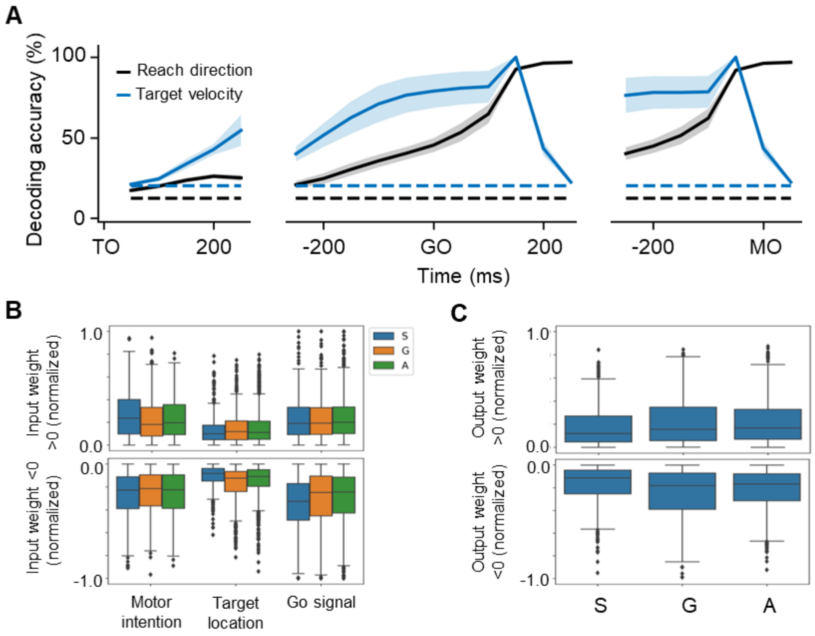


**Figure S12. Decoding results and weight distributions of RNNs**

**A** The decoding accuracy (SVM with 10-fold cross-validation) of reach direction (black line) and target velocity (blue line) across 100 network models, is aligned to the TO, GO, and MO. The dash-dotted lines are chance level of decoding reach direction (black, one in eight) and that of target velocity (blue, one in five). The lines and the shaded area indicate the mean and the standard deviation across models, respectively.

**B** The input weights. The weights from motor intention-x and motor intention-y were averaged to obtain “motor intention”, so as target location-x and target location-y to “target location”. To view relative tendencies, we selected solely modulated nodes, and normalized the weight by dividing it by the corresponding maximum (the max absolute weight from motor intention, target location, and GO-signal, respectively) for each model. For motor intention, S>G=A (weight>0); for target location, S<G=A (weight >0), S<A<G (weight<0); for GO-signal, S>G=A (weight<0). ‘=’ here defines no significance while ‘<’ means p<0.01 (K-W test).

**C** The output weights. The outputs to x and y were averaged. Similar to **C**, we selected the solely modulated nodes and normalized the weights by the max absolute output weights, for each model regardless of whether weight >0, S<G=A (K-W test, p<0.001).
